# TurboID-mediated surface protein biotinylation to inhibit the growth of *Staphylococcus aureus*

**DOI:** 10.1101/2024.06.24.600537

**Authors:** Lijuan Qian, Yuxin He, Wenzhe Lian, Zhiyuan Ji, Ziming Tian, Chuyun Wang, Chen Cao, Tyler Shern, Teagan Stedman, Yujun Sun

## Abstract

*Staphylococcus aureus* (*S. aureus*) infection is major cause of nosocomial infections. Antibiotic treatment for *S. aureus* remains the primary solution for managing *S. aureus* infections, which, however, increases the risk of antibiotic resistance. To broaden the resolutions on *S. aureus* infection, here we report TurboID-mediated protein proximity technologies to inhibit the growth of *S. aureus*. To achieve this goal, we utilized synthetic biology techniques to create a fusion protein named N-AgrD-TurboID (Agr-ID). The N-AgrD domain includes auto-inducer peptide (AIP) which combined to the surface AgrC protein on *S. aureus*. As such, TurboID then catalyzed the production of biotinoyl-5’-AMP anhydride, triggering the biotinylation of surface proteins on *S. aureus* 25923 which were visualized by using fluorescence microscopy after incubating with Alexa Fluor 647-conjugated streptavidin. The biotinylation of surface protein on *S. aureus* 25923, *S. aureus* 43300, and *S. aureus* 6538 (MRSA) also resulted in growth inhibition and impaired colonization. Moreover, the biotinylation on surface protein further inhibited virulence protein production in *S. aureus* 25923, as indicated by reduced apoptosis of HEK 293T cells after treatment with *S. aureus* 25923 lysates. Overall, our work reveals that the biotinylation of surface proteins can inhibit the growth and toxicity of *S. aureus* 25923, *S. aureus* 43300, and *S. aureus* 6538 (MRSA), indicating therapeutic potential in clinical treatment.

## Introduction

*Staphylococcus aureus* (*S. aureus*) is a pathogenic bacterium that can cause multiple fatal infections, including skin, respiratory, and blood infections, and was the leading cause of death in the United States (Dayan et al., 2016). Over the past decade, *S. aureus* has been reported to cause 50 cases per 100,000 annually with a mortality rate of 30% (Cheung et al., 2021). The high mortality rate is mainly attributed to the high drug resistance of *S. aureus*. Currently, only the clinical drugs vancomycin and rifampicin have been approved for use in the treatment of methicillin-resistant *S. aureus* (MRSA) infections (Kumar, 2016). However, the evolution of vancomycin resistance has accelerated with its widespread use (Stogios and Savchenko, 2020). Other, glycopeptide drugs, such as dalbavancin and telavancin, are alternative ways to treat *S. aureus* infection (Roecker and Pope, 2008). Inhibition of enzymes related to cholesterol biosynthesis can be an effective way to detoxify *S. aureus* (Liu et al., 2008). 1,4-dihydroxyphosphonic acid inhibits the sa-Men D enzyme, thereby inhibiting the bacterial respiratory chain of *S. aureus* (Stanborough et al., 2023). However, treatment methods against *S. aureus* infections are still limited.

Quorum sensing (QS) is a major driver of virulence and antibiotic resistance, making it a promising system to target therapeutically. Notably, the inhibition of surface receptor proteins shows great promise in combating *S. aureus* infections. Surface Agr inhibitors can directly block the quorum sensing signaling pathway of *S. aureus*, such as inhibiting the autoinducer peptide-sensing protein, AgrC (Kaur et al., 2021; Otto, 2023). Inhibiting the activity of surface Agr leads to a reduction in the expression of virulence genes in *S. aureus* (Gordon et al., 2013). AgrC-mediated biofilm formation is a crucial factor in the pathogenicity of *S. aureus* (Chan et al., 2004). Notably, planktonic *S. aureus* is 10 to 1000 times more susceptible to antibiotics than biofilm-associated strains (Winkler and Haiden, 2017). The production of biofilms relies on the activation of AgrC by the autoinducer peptide, which then phosphorylates AgrA, inducing RNAIII production, and promoting biofilm dissociation (Li and Tian, 2012). The inhibition of AgrC using synthetic epidermis pheromones or a membrane-embedded peptidase suppresses toxin production and biofilm colonization by *S. aureus* (Cosgriff et al., 2019; Vuong et al., 2000).

Thus, inhibiting surface Agr can further suppress toxin and biofilm production. Protein modification, such as biotinylation, offers an efficient approach to inhibit surface Agr function (Horswill and Gordon, 2020). TurboID serves as a tool that enables the efficient biotinylation of binding partners of a protein of interest through the fusion of a promiscuous biotin ligase (Branon et al., 2018). Leveraging TurboID, we developed a fusion protein, N-AgrD-AIP-TurboID (Agr-ID) containing two domains: N-AgrD-AIP and TurboID. The specific interaction between N-AgrD-AIP and surface AgrC proteins can bring TurboID closer to surface AgrC. Biotinylation of surface proteins further impedes biofilm production and the expression of virulence genes, ultimately inhibiting the growth and survival of *S. aureus*.

## Materials and Methods

### Materials

The *S. aureus* strains ATCC25923, *S. aureus* ATCC43300, and *S. aureus* ATCC6538 (MRSA) was purchased from China Industrial Microbiological Culture Collection and Management Center. *Escherichia coli* (ATCC25922), *Pseudomonas aeruginosa* (ATCC27853) and *Bacillus sp*. CZGRY11 were provided by Horticulture Laboratory of Anhui Science and Technology University. Yeast powder was purchased from Oxoid Ltd; Peptone was purchased from Beijing Aoboxing Biotechnology Co., Ltd; Imidazole and sodium chloride were purchased from National Pharmaceutical Group Chemical Reagents Co., Ltd; Tris-HCl and agar powder were purchased from Beijing Solebold Technology Co., Ltd; Polyacrylamide gel was purchased from Nanjing Novozan Biotechnology Co., Ltd; Chloramphenicol and Isopropyl β-D-1-thiogalactopyranoside (IPTG) were purchased from Beyotime Biotech Inc; The Nickel column filler was purchased from Kingsley Biotechnology Co., Ltd.

### Plasmid construction

The TurboID-V5 gene sequence was obtained from AddGene (#107167). The AgrD_1-32_ domain consists of an N-terminal domain from amino acid 1-27 and the AIP region from amino acid 28-32. The fusion gene was generated using SnapGene software and synthesized by Beijing Genomics Institute (BGI). The fusion gene was cloned into a pET-22b plasmid for protein expression and purification.

### Protein purification

The recombinant plasmid was transformed into BL21 (DE3) competent cells and plated on solid medium containing 100 μg/mL ampicillin. After 18 h, single colonies were selected for Sanger sequencing as part of the quality control (QC) process. The strains that passed the QC test were then cultured in 200 mL of Lysogeny Broth (LB) media at 37 °C and 200 rpm for approximately 4-6 h. After reaching an optical density at 600 nm (OD_600_) of 0.6, 200 µL of 1M Isopropyl β-D-1-thiogalactopyranoside (IPTG) was added to the medium, followed by continuous culturing at 16 °C for an additional 8 ho. Finally, the supernatant was removed after centrifugation at 4 °C and 8000 rpm for 10 min. The pellet was then washed three times with 10 mL of 1x PBS before proceeding to lysis. Subsequently, 10 mL of a 5 mM imidazole solution was added to the collected bacteria, which were then subjected to ultrasonic crushing for 20 min using cycles of 9 s on and 6 s off. Bacterial lysates were then centrifuged at 4 °C and 10,000 rpm for 15 min. The QC for protein expression was assessed by conducting 10% SDS-PAGE followed by Coomassie staining. Ni^+^ column was pre-equilibrated by adding 10 mL of 5 mM imidazole and washing it 3 times. Before passing through the Ni_+_ column, the supernatants were filtered using a 0.45 μm filter membrane. Ni^+^ based protein purification methods were followed by the protocol from Genscript.

### The biotinylation reaction on *S. aureus* strains and visualization

To address the questions, Agr-ID gene were designed and cloned into a pET-22b plasmid. In consideration of protein folding and stability of Agr-ID, a pelB signal peptide was used to translocate the Agr-ID to periplasm (Kusuma et al., 2022; Singh et al., 2013). The Agr-ID protein was purified using a Nickel (Ni) column affinity chromatography method, facilitated by the presence of a 6x Histidine (His) tag on the protein. The successful purification of Agr-ID was confirmed by observing the expected band corresponding to Agr-ID on a 10% SDS-PAGE gel after Coomassie staining.

The 100 µL reaction system contained 1 µL *S. aureus* (∼1.66 x 10^9^), 5 µL 100 mM ATP, 5 µL 5 mM Biotin, 5 µL 500 mM MgCl_2_, 34 µL Buffer A (2.5 mL 1 M Tris-HCl pH=8.0, 6.25 mL 4 M NaCl, 1.25 mL 2 M Imidazole), 50 µL 0.24 mg/mL Agr-ID purified protein. The reaction was performed at 30°C for 12 h, followed by culturing on LB solid medium or in liquid LB medium to assess the effects on bacterial growth.

Next, the biotinylating reaction was performed in the 100 µL buffer containing 5 µg Agr-ID protein and 1.6 x 10^9^ *S. aureus* cells. The reaction is terminated by adding 3 mL Tris-buffer and collecting bacteria at 8000x g for 15 min. The biotinylated *S. aureus* cells were then stained by incubating 1 mM the Alexa Fluor 647-conjugated streptavidin and visualized fluorescent microscope.

### Bacterial growth curves

All the biotinylated *S. aureus* from the previous step were cultured in LB medium with or without 0.24 mg/mL Agr-ID purified protein, 5 µL 100 mM ATP, 5 µL 5 mM biotin. The growth of *S. aureus* was tracked by measuring OD_600_ at 0, 2, 4, 8, and 18 h time points.

### Cytotoxic detection by using Annexin V

The virulence protein produced by *S. aureus* were measured by testing the apoptosis of co-cultured human HEK 293T cells (Wesson et al., 2000). In brief, HEK 293T cells were cultured in RPMI media with 10% FBS and harvested using 0.25% trypsin-EDTA. 1 million cells were resuspended with 1 mL 10% FBS RPMI media. For making the bacterial lysate, *S. aureus* were cultured in LB media and harvested at an OD_600_ of 1.0 (approximately 3 × 10^7^ cells/mL). 1 mL *S. aureus* were centrifuged at 12,000x g for 5 min and were resuspended in 100 µL RPMI media. *S. aureus* were sonicated for 5 min and centrifuged at 12,000x g for 5 min. The supernatant was then added to the 293T cells for 30 min in 37 °C.

After the reaction, harvest the cells by centrifuging at 400x g for 5 min. The cells were resuspended in 100 µL 1x Annexin V binding buffer and stained with Annexin V-FITC for 15 min and then DAPI staining for 5 min. The staining was ended by adding with 1 mL 1x Annexin binding buffer and centrifuged at 400x g for 5 min. Cell were resuspended in 100 µL 1x annexin binding buffer and evaluated in flow cytometry.

### Statistical analysis

Experiments were statistically analyzed using Graph Pad Prism8. The normality test was first performed using Shapiro-Wilk test (n < 8). Data that passed the normality test were undergoing the two-tailed Students’ *t*-test for the comparison of two groups or two-way analysis of variance (ANOVA) with Tukey’s multiple comparisons. The data that passed the normality test were represented by the mean ± standard error of the mean (SEM). Two-sided P values were determined by unpaired student t test for comparing data that did not pass the normality test. Data were then presented as median ± 95% confidence interval (CI). Statistical significance of difference was accepted when *P* values were < 0.05. All the specific *P* values and biological replicants were listed in figures and figure legends.

## Results

### The construction and characterization of Agr-ID

Novel methods for inhibiting the quorum sensing (QS) sensor protein surface Agr have demonstrated effective promise in detoxifying *S. aureus* (Kaur et al., 2021; Otto, 2023), indicating surface Agr protein has been an important potential therapeutic target. It has been established that post-translational protein modifications are a crucial strategy contributing to the disruption of protein function, including protein biotinylation (Rodriguez-Melendez and Zempleni, 2003) Here, we hypothesis a novel method for the inhibition of the QS receptor surface Agr, we employed the TurboID protein to catalyze surface Agr biotinylation. As shown in Fig.1A, a fusion expression approach was employed, linking AgrD to the C-terminus of TurboID. This resultant fusion protein was designated as Agr-ID. The N-terminal domain of Agr-ID contains an autoinducer peptide (AIP) aiming at interacting with AgrC (Bardelang et al., 2023; Xiang et al., 2020). The N-terminal Agr-ID domain functioned as a “finding” domain to localize Agr-ID in close proximity to the surface of *S. aureus*, whereas the C-terminal TurboID domain acted as a “catalytic” domain. TurboID facilitated the biotinylation of all exposed lysine residues within a 10 nm radius by the effective conversion of ATP and biotin to form a reactive biotinoy1-5’-AMP conjugate. Thus, two subsequent questions must be addressed: 1) whether the biotinylation of surface protein including AgrC on *S. aureus* can be achieved. 2) what changes happened to biotinylated *S. aureus*.

### Agr-ID inhibited the growth of *S. aureus* 25923, *S. aureus* 43300, and *S. aureus* 6538 (MRSA)

After performing the biotinylating reaction, the biotinylated *S. aureus* were successfully visualized by using Alexa Fluor 647-conjugated streptavidin and fluorescent microscope. By utilizing the total cell lysates with unpurified Agr-ID,around 10% of total 1.6 x 10^9^ *S. aureus* cells were labeled biotin indicating a feasible TurboID-based protein proximity labeling method for the treatment of *S. aureus* (Fig.1B). To further investigate the growth inhibition of biotinylated *S. aureus*, we then firstly purified and condensed Agr-ID before adding to the biotinylating reaction buffer. This purified method were increased nearly 5 folded efficiency of biotinylation on *S. aureus* (Fig.1C). We then employed the growth curve measurement by detecting OD_600_ which identified as a most commonly used way to monitor the growth of bacteria (Krishnamurthi et al., 2021). As shown in (Fig. 1D), there was no significant difference in the first 2 h, indicating that the biotinylated *S. aureus 25932* were not dying in the first 2 h. However, the growth inhibition showed significant difference between the two groups suggesting that the blocking of surface AgrC protein by Agr-ID caused growth arrest after 2 h blocking by surface Agr protein biotinylation. More importantly, as known that biofilm-associated genes are usually activated when the OD_600_ reaches 0.8 (Wu et al., 2015). We observed that the growth of biotinylated *S. aureus 25932* did not reach an OD_600_ value of 0.8, indicating reduced quorum sensing (QS) signaling and inhibited biofilm production in this strain. To further confirmed the growth inhibition of biotinylating on other strains of *S. aureus*, we employed two more strains, including *S. aureus* 6538 (Fig. 1E) and *S. aureus* 43300 (Fig. 1F). Under biotinylating condition, both *S. aureus* strains showed growth arrest which indicates the success of biotinylating modification on the surface AgrC proteins of *S. aureus* 43300 and leads to the potential use of this methods for anti-bacterial treatments.

**Figure 1.**
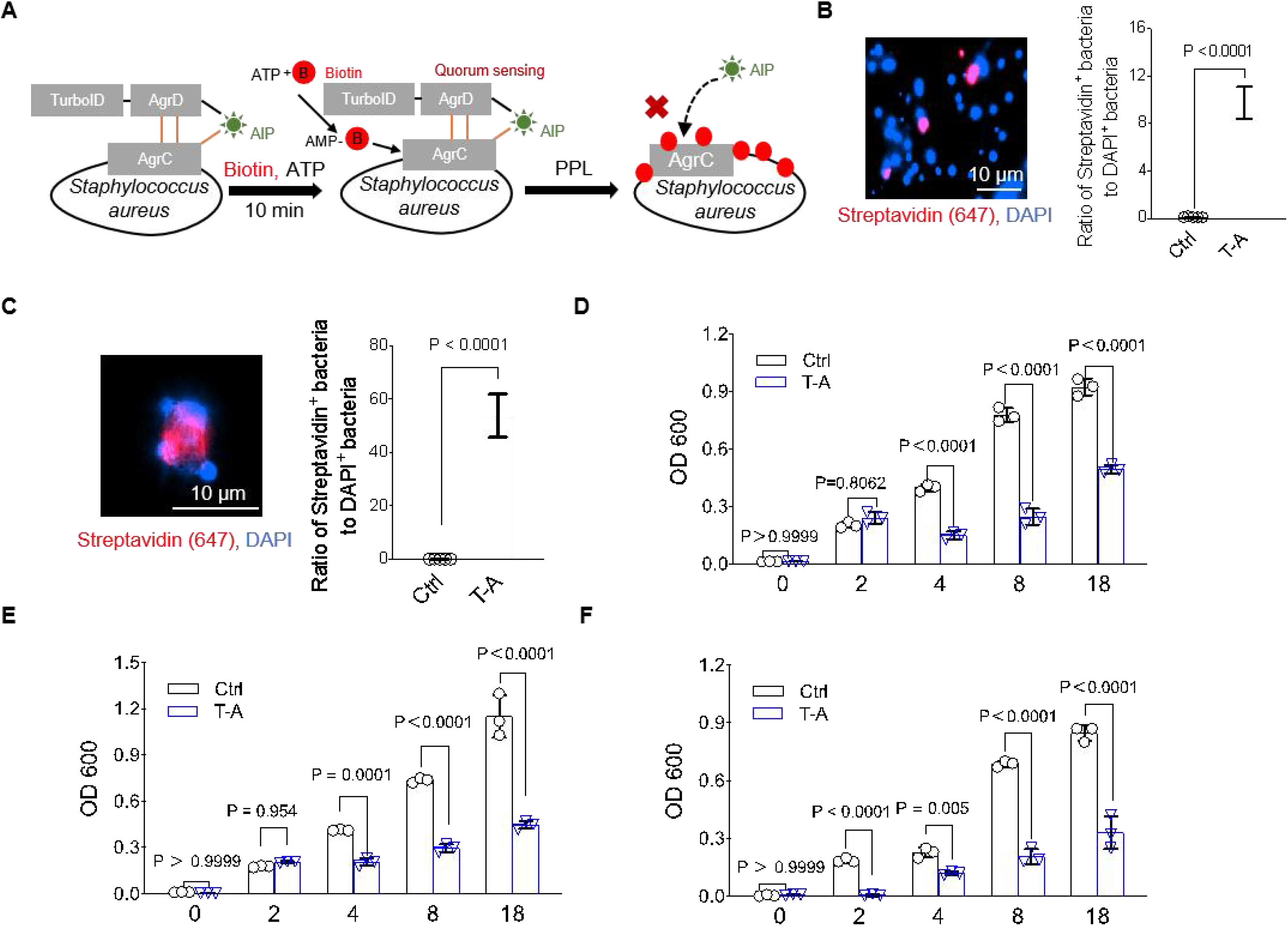
TurboID-base protein proximity biotinylation on *Staphylococcus aureus* surface protein and its inhibition on growth. **(A)** The N-terminal domain of AgrD gene were fused with TurboID to further generate the fusion protein Agr-ID. The N-ternimal AgrD domain contained auto-inducer (AIP) domain which guided TurboID getting closer to AgrC, the main sensor proteins for quorum sensing (QS). After adding ATP and Botin in the reaction buffer, TurboID catalased ATP and Biotin to format AMP-biotin which further biotinylated the lysine residues on Agr. The biotinylation occurring on Agr will blocked its function in sensing AIP in the environment. Consequently, growth is inhibited, and endotoxic genes are silenced in *S. aureus*. **(B, C)** Growth inhibition in biotinylated *S. aureus 25923*. The percentage of biotinylated *S. aureus* was qualified by calculating the ratio of Streptavidin+ *S. aureus* versus total DAPI^+^ *S. aureus*. Data are presented as mean ± SEM. Two-sided *P* values were determined by unpaired student t test. N=5 independent biological replicants. **(D-F)** Growth inhibition observed after surface biotinylating modification of *S. aureus* 25923 (D), 6538 (E), 43300 (F). Non-biotinylated and biotinylated *S. aureus* were cultured in LB media culturing and tracking for indicated 0, 2, 4, 8, 18 h. OD_600_ value were collected and calculated from 3 independent biological replicants. N=3 independent biological replicants. Data are presented as mean ± SEM. Two-sided *P* values were determined by a two-way ANOVA with Tukey’s multiple comparisons test. T-A: Agr-ID.

### Agr-ID inhibited the colonization of *S. aureus* 25923

Colonization of bacteria was recognized as the most essential and primary step before the growing and multiplying from host body. To further investigate the inhibition of colonization on S. aureus, we employed three strains including antibiotic resistant stain *S. aureus* 6538 (MRSA). The number of colonies in the group of biotinylated *S. aureus* 25923 (Fig. 2A) were significantly reduced, indicating the surface protein biotinylation affected the conization directly within the first 2 h.

**Figure 2.**
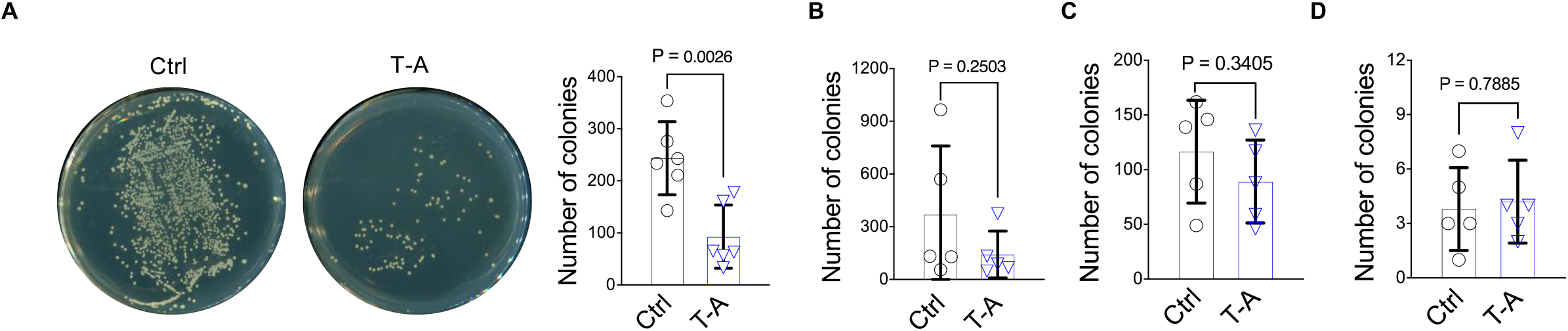
Biotinylated *S. aureus* 25923 showed specifically decreased colonization, not on others species. **(A)** Inhibited colonization of three biotinylated *S. aureus* strains including *S. aureus* 25923. Data are presented as mean ± SEM. Two-sided *P* values were determined by unpaired student t test. N=4-6 independent biological replicants as indicated. **(B-D)** Biotinylating-mediated colonization inhibition of three none *S. aureus* strains after reaction including *Escherichia coli* (B), *Pseudomonas aeruginosa* (C) and *Bacillus subtilis* (D). N=4-5 independent biological replicants as indicated.

Moreover, to investigate Agr-ID in recognition and specificity on other species with no surface Agr proteins. We employed *Escherichia coli, Pseudomonas aeruginosa* and *Bacillus sp*. CZGRY11 and performed with the same colonization assay. As shown in Fig. 2B-D, Agr-ID was not successful in inhibiting the growth of the three indicated bacterial strains, suggesting that if “finding” AgrD domain can not successfully recognized the other species with non-Agr expressed, the “catalytic” TurboID domain can not work properly to get the bacterial surface protein biotinylated.

### Agr-ID reduced the virulence of *S. aureus* 25923

To further investigate the production of virulence proteins in *S. aureus* 25923, lysates of *S. aureus* 25923 were utilized to assess their ability to induce apoptosis in mammalian cells (Wesson et al., 1998). After 18 h of culture, equal numbers of non-biotinylated and biotinylated *S. aureus* 25923 were harvested and then lysed using the sonication method. The supernatant was then added to the culture media of HEK 293T cells and co-cultured for 3 h, followed by harvesting the cells and staining with Annexin V/DAPI. As depicted in (Fig. 2A), the population of healthy HEK293T cells significantly increased (Fig. 2B), whereas the rate of early apoptosis showed no significant change (Fig. 2C). Meanwhile, the number of late apoptotic (Annexin V^+^DAPI^+^) HEK293T cells decreased (Fig. 2D). These results reflected the the biotinylated *S. aureus* 25923 were less toxic compared with non-biotinylated *S. aureus* 25923.

## Discussion

*S. aureus* infection has been significant global health challenge (Kadariya et al., 2014). Antibiotic treatment has come a long way in addressing *S. aureus* infections. However, long-term use of antibiotics causes antibiotic resistance (Salisbury et al., 2002), increased susceptibility to infections (Lowy, 1998), and adverse effects on overall immune function (Langdon et al., 2016). Therefore, a novel protective method that can specifically inhibit the virulence of *S. aureus* is needed. Activation of the QS system, necessary for the expression of *S. aureus* virulence genes, is largely dependent on the accumulation of autoinducer (AIP) in the environment (Wang and Muir, 2016). The QS system continuously produces AIP, eventually activating itself through the AIP and surface Agr protein interaction (Scoffone et al., 2019). Recent advancements in inhibiting the quorum sensing (QS) sensor protein surface Agr have shown promising effectiveness in detoxifying *S. aureus* (Kaur et al., 2021; Otto, 2023). Hence, we developed the fusion protein Agr-ID with the aim of targeting the loss of function of the surface Agr protein through biotinylation.

In this study, we initially assessed the potential for loss of function in the surface Agr protein of *S. aureus* through the specific biotinylation of lysine residues using Agr-ID. However, the surface protein on *S. aureus* should be pulled-down to further confirm the success in biotinylation modification on AgrC and also the potential transient and covalent interactomes with AgrC on the surface of *S. aureus*. These will further confirm the growth inhibition and impaired colonization were regulating by the loss-of-function of AgrC or other proteins.

Moreover, the virulence genes downstream of AgrC need to be evaluated to further explained the causal genes for the apoptosis of HEK293T cells in Fig.3 by virulence protein in *S. aureus* 25923 cell lysates. Thirdly, the potential clinic application of Agr-ID will be evaluated. In healthy adults, the serum concentration of ATP is approximately 1 mM (Gorman et al., 2007), while the range of biotin concentration is from 133-329 pM (Sweetman and Nyhan, 1986). This is an idea system for Agr-ID to leverage the *S. aureus* infection. As such, Pre-clinic studies are needed to test the feasibility of Agr-ID in anti-*S. aureus* infection.

**Figure 3.**
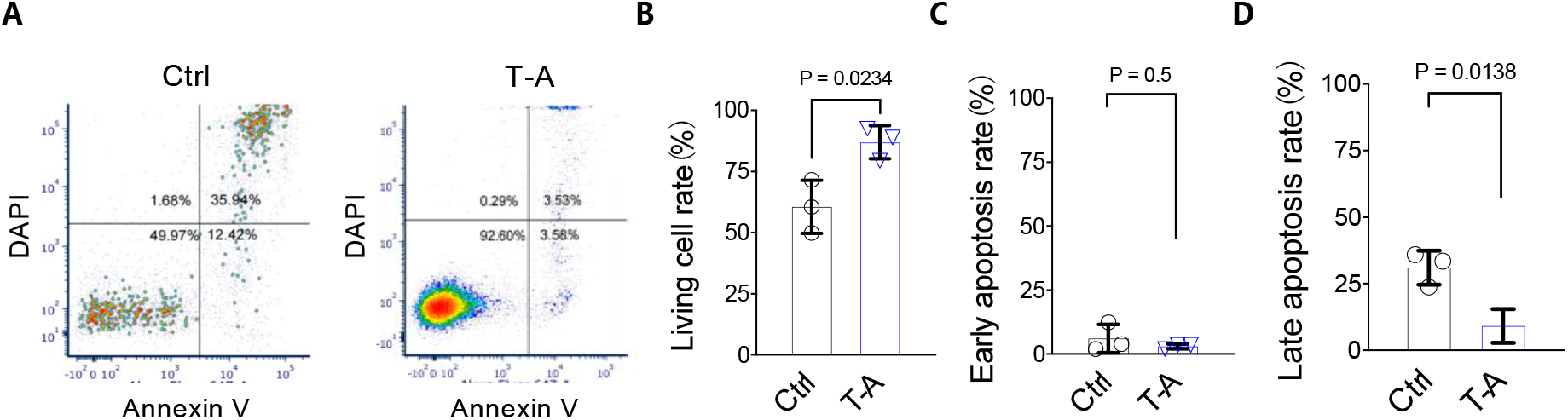
The inhibition of virulence genes expression in biotinylated *S. aureus* 25923. **(A)** Non-biotinylated and biotinylated *S. aureus* 25923 were cultured and lysed by using ultrasonicator. After centrifuging, the supernatants were applied to 293T cells and incubated for 3 hours. 293T cells were harvested and performed with Annexin V staining and DAPI staining for 10 mins and detected by flow cytometry. representative figures for indication of gating strategy for testing cell apoptosis. **(B-D)** calculations on live cell rate **(B)**, early apoptosis **(C)**, and late apoptosis **(D)**. Data were collected and calculated from 3 independent biological replicants. Data are presented as mean ± SEM. Two-sided *P* values were determined by unpaired student t test. N=3 indicated as three independent biological replicants. T-A: Agr-ID.

## Conclusion

Agr-ID demonstrates enhanced specificity and efficiency in biotinylating the surface AgrC protein on *S. aureus* 25923. This presents a promising alternative avenue for addressing *S. aureus* 25923, *S. aureus* 43300, and *S. aureus* 6538 infections. Biotinylation of surface AgrC results in the loss of function in sensing the quorum sensing (QS) signaling peptide AIP, leading to the inhibition of virulence gene expression. This study introduces a novel method for inhibiting QS signaling through protein biotinylation modification. The techniques developed here have the potential for broader application with inhibition of other pathogenic bacteria.

## Author Contribution

LJQ and YJS: survey, writing - original, writing - review, editing, conceptualization, methodology, resources, data management, investigation, project management, validation, visualization, writing - manuscript, writing - review and editing, software, supervision, project administration. LJQ and YXH: methodology, conceptualization, writing - review and editing. WZL, ZYJ, ZMT, CYW, CC, Tyler S, Teagan S: methodology, formal analysis and writing review.

## Full Data Availability

All authors declare that the data supporting the findings of this study are available within the article.

